# *In planta* expression screens of candidate effector proteins from the wheat yellow rust fungus reveal processing bodies as a pathogen-targeted plant cell compartment

**DOI:** 10.1101/032276

**Authors:** Benjamin Petre, Diane G.O. Saunders, Jan Sklenar, Cécile Lorrain, Ksenia V. Krasileva, Joe Win, Sébastien Duplessis, Sophien Kamoun

## Abstract

Rust fungal pathogens of wheat (*Triticum* spp.) affect crop yields worldwide. The molecular mechanisms underlying the virulence of these pathogens remain elusive, due to the limited availability of suitable molecular genetic research tools. Notably, the inability to perform high-throughput analyses of candidate virulence proteins (also known as effectors) impairs progress. We previously established a pipeline for the fast-forward screens of rust fungal effectors in the model plant *Nicotiana benthamiana*. This pipeline involves selecting candidate effectors *in silico* and performing cell biology and protein-protein interaction assays *in planta* to gain insight into the putative functions of candidate effectors. In this study, we used this pipeline to identify and characterize sixteen candidate effectors from the wheat yellow rust fungal pathogen *Puccinia striiformis* f sp *tritici*. Nine candidate effectors targeted a specific plant subcellular compartment or protein complex, providing valuable information on their putative functions in plant cells. One candidate effector, PST02549, accumulated in processing bodies (P-bodies), protein complexes involved in mRNA decapping, degradation, and storage. PST02549 also associates with the P-body-resident ENHANCER OF mRNA DECAPPING PROTEIN 4 (EDC4) from *N. benthamiana* and wheat. Our work identifies P-bodies as a novel plant cell compartment targeted by pathogen effectors.

## INTRODUCTION

Plant pathogens colonize hosts by deploying virulence proteins known as effectors that manipulate plant cell structures and functions (Dodds and Rathjen, 2010; Win *et al*., 2012). Once delivered into host tissues, effectors reside in the extracellular space (apoplastic effectors) or translocate into the plant cells (cytoplasmic effectors). Unravelling how effectors function in the host is key to understanding parasitism and to developing resistant plants (Dangl *et al*., 2013). Pathogen effectors are operationally plant proteins; they function in plant tissues, they associate with plant molecules, and their phenotypic expression in plants drives their evolution (Hogenhout *et al*., 2009). The field of effector biology has rapidly advanced in recent years due in large part to the availability of host plants that are amenable to molecular genetics and in which effectors can be heterologously expressed and studied (for comprehensive reviews, see Martin and Kamoun, 2012). However, due to the limited availability of functional genetic resources for crop species, characterising the effectors of crop pathogens remains challenging (Petre *et al*., 2014; Upadhyaya *et al*., 2014). To study crop pathogen effectors, an alternative approach is the use a surrogate experimental plant system, such as *Nicotiana benthamiana* (Petre *et al*., 2015a).

*Nicotiana benthamiana* (Solanaceae) is a well-established experimental system to study proteins *in planta* (Goodin *et al*., 2008; Bombarely *et al*., 2012). The agroinfiltration method allows transient expression of proteins in leaf cells, and a wide range of assays is available for functional investigations. Thus, *N. benthamiana* is extensively used in effector biology (Pais *et al*., 2014). We recently used this plant to set up an effectoromics pipeline aimed at determining the plant cell compartments and protein complexes targeted by candidate effectors of rust fungi (Petre *et al*., 2015a; Petre *et al*., 2015b). Such pipeline is a valuable tool for the rapid screening of candidate effectors.

Rust fungi (Pucciniales) are notorious for being destructive crop pathogens (Dean *et al*., 2012). The species that infect wheat pose a constant threat to global food security (Beddow *et al*., 2015). These fungal pathogens include the yellow rust fungus *Puccinia striiformis* f sp *tritici* (Chen *et al*., 2014; Hubbard *et al*., 2015). To date, effectors have not been functionally characterized for this species. However, genome and transcriptome analyses have predicted hundreds of candidate effectors, most of which are secreted proteins of unknown function (Cantu *et al*., 2011; Garnica *et al*., 2013; Zheng *et al*., 2013). Cantu and colleagues recently combined genome and *in silico* analyses to prioritize candidate effectors for further functional analyses (Cantu *et al*., 2013).

Processing bodies (P-bodies) are protein/RNA complexes that reside in the cytosol of eukaryotic cells. They control the decapping, degradation, and storage of mRNA molecules (Chan and Fritzlers, 2012). In plants, P-bodies and P-body-resident proteins have important roles in post-embryonic development (Xu *et al*., 2006; Xu and Chua, 2009), salt stress tolerance (Steffens *et al*., 2015), and immune responses (Maldonado-Bonilla *et al*., 2014). In animals and yeast, limited evidence suggests that pathogenic bacteria and viruses target P-bodies (Eulalio *et al*., 2011; Reineke and Lloyd, 2013). To date, no connection has been made between P-bodies and filamentous pathogens.

In this study, we investigated sixteen candidate effectors of the wheat yellow rust fungus *P. striiformis* f sp *tritici* using the *N. benthamiana* effectoromics pipeline we previously developed (Petre *et al*., 2015a). We discovered that nine candidate effectors accumulate in distinct plant cell compartments and associate with specific protein complexes. Notably, the candidate effector PST02549 accumulates in P-bodies and associates with the wheat enhancer of mRNA decapping protein 4. Our findings suggest that P-bodies are a plant compartment targeted by pathogen effectors. We also conclude that *N. benthamiana* can be used as an experimental system to screen candidate effectors of pathogens of monocot plants, including obligate biotrophic pathogens of wheat.

## RESULTS

### Selection of 16 candidate effectors from *Puccinia striiformis* f sp *tritici*

The predicted effector complement of *P. striiformis* f sp *tritici* consists of hundreds of secreted proteins (Cantu *et al*., 2013). To select candidate effectors for functional investigations, we leveraged our recently developed pipeline (Petre *et al*., 2015a) to select eleven proteins, using transcript enrichment in purified haustoria as the principal criterion for selection. We also included five proteins previously flagged as promising candidates (Cantu *et al*., 2013) to obtain a final list of sixteen candidate effectors (Table 1, Table S1). These sixteen candidates are Pucciniales-specific, and only seven show some sequence similarity to proteins of the wheat stem rust fungus *Puccinia graminis* f sp *tritici* (Table 1). This finding suggests that most of these candidate effectors recently emerged in the Pucciniaceae family.

**Table 1.** *Puccinia striiformis* f. sp. *tritici* candidate effectors analysed in this study. ^a^ Protein IDs were adapted from Cantu *et al*., 2013 by removing the isolate ID for simplicity. Full-length protein IDs are indicated in Table S1. ^b^ Data mined from Cantu *et al*., 2013. ^c^ Number of amino acids are indicated; SP: predicted signal peptide ^d^ Number of cysteine residues in the mature form of the protein (i.e. without the signal peptide) ^e^ H: transcripts were identified in isolated haustoria; H5: transcripts were in the top 5% of transcripts detected in isolated haustoria; H1: transcripts were in the top 1% transcripts detected in isolated haustoria; I: transcripts were identified in wheat tissues during infection; I10: transcripts were in the top 10% transcripts detected in wheat tissues during infection; TC: transcript accumulation was detected by RTq-PCR during a time-course infection of wheat. ^f^ Pgt: *Puccinia graminis* f sp *tritici;* Mlp: *Melampsora larici-populina;* Hv: *Hemileia vastatrix*.

### Candidate effector-fluorescent protein fusions accumulate in *N. benthamiana* leaf cells

To test whether the mature form (i.e. without the signal peptide) of the candidate effectors could be expressed in dicot cells, we generated candidate effector-green fluorescent protein (GFP) fusions and expressed them in *N. benthamiana* by agroinfiltration. Live-cell imaging and immunoblotting assays revealed that the sixteen fusion proteins accumulate in leaf cells at detectable levels, with no obvious sign of aggregation or degradation (Figure 1; Figure S1). Some fusions showed a band signal at a lower molecular weight, in addition to the band signal at the theoretical size (PST18220, PST03196, and PST12160), or a band signal at a higher molecular weight than expected (PST02549 and PST05023), suggesting post-translational modifications (Figure S1). As the proteins effectively accumulated in leaf cells, we inferred that transient assays in *N. benthamiana* are suitable for further *in planta* analyses.

**Figure 1.**
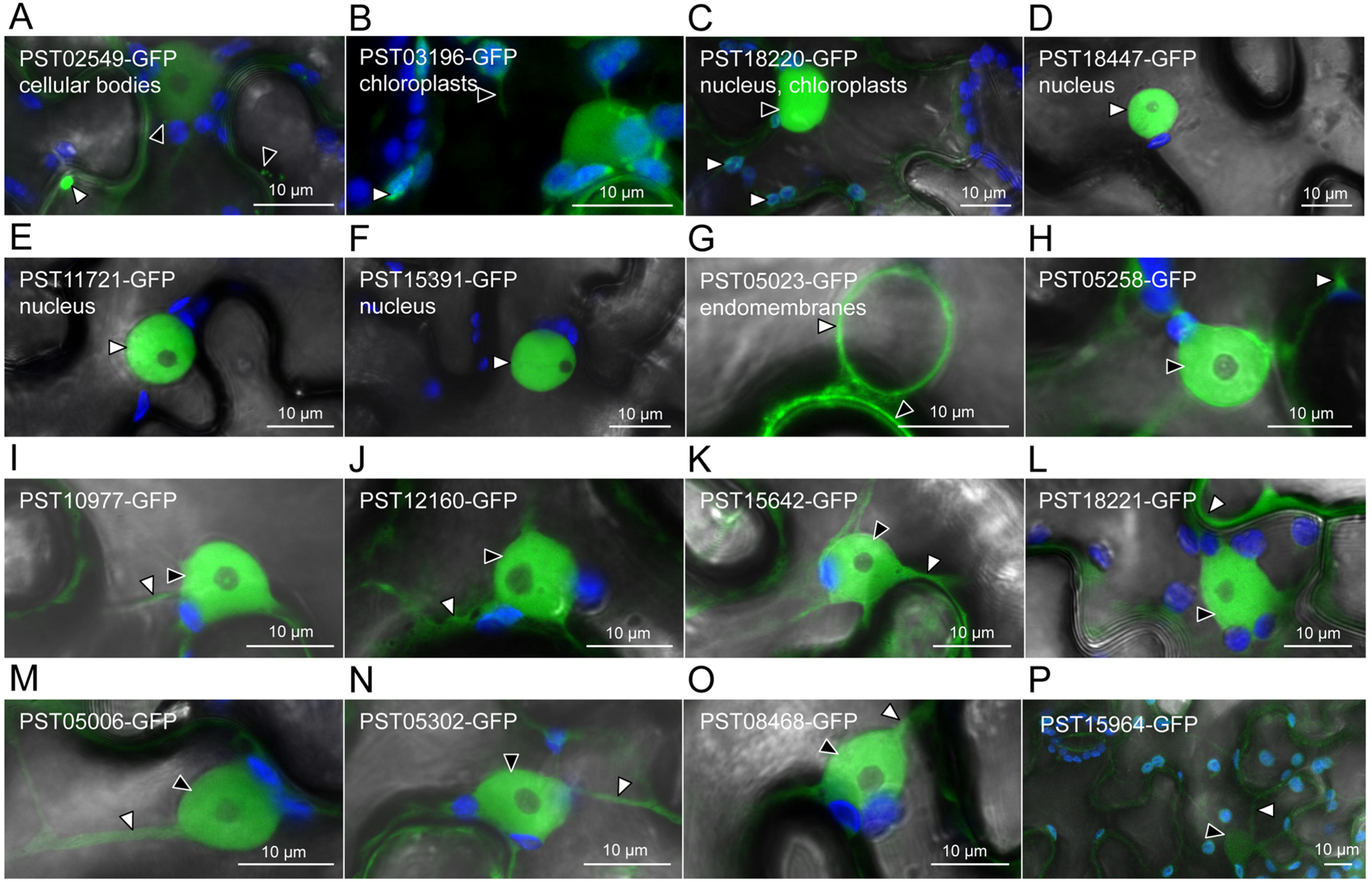
Seven candidate effectors show specific accumulation patterns in leaf cells. Live-cell imaging of the 16 candidate effector-GFP fusion proteins accumulating in distinct subcellular compartments of *N. benthamiana* leaf cells. Proteins were transiently expressed in *N. benthamiana* leaf cells by agroinfiltration. Live-cell imaging was performed with a laser-scanning confocal microscope two days after infiltration. GFP and chlorophyll were excited at 488 nm. GFP (green) and chlorophyll (blue) fluorescence were collected at 505-525 nm and 680-700 nm, respectively. Images are single optical sections of 0.8 µm or a maximal projection of up to 47 optical sections (max. z-stack of 37.6 µm). Images displayed are overlays of the GFP signal, the chlorophyll signal, and bright field. For A-G, specific cellular compartments in which the GFP signal accumulates are indicated. White arrowheads indicate GFP-labelled cytosolic bodies (A), chloroplasts (B-C), nuclei (D-F), nuclear surrounding (G), or cytosolic fractions (H-P). Black arrowheads indicate GFP-labelled small cytosolic bodies (A), a stromule (B), a nucleus (C), the plasma membrane (G), or nuclei (H-P). In (P), the low level of accumulation of the fusion protein imposed higher laser power and gain, which resulted in non-specific signal for the GFP channel being visible in chloroplasts and ostiole edges.

### Seven candidate effectors accumulate in specific plant cell compartments

To identify the plant cell compartments in which the candidate effectors accumulate, we performed live-cell imaging of cells expressing effector-GFP fusions. Seven out of the sixteen fusion proteins displayed an informative distribution in leaf cells (Figure 1). The fluorescence signal from PST02549-GFP and PST03196-GFP accumulated in small cytosolic bodies and chloroplasts, respectively, as well as in the nucleus and the cytosol (Figure 1A-B). The fluorescence signal from PST18220-GFP labelled both chloroplasts and nuclei, suggesting a dual targeting to the two organelles (Figure 1C). The fluorescence signal from PST18447-GFP, PST11721-GFP, and PST15391-GFP specifically accumulated in nuclei, and PST11721-GFP also labelled nuclear foci in some rare cases (Figure 1D-F, Figure S2). Finally, the fluorescence signal from PST05023-GFP labelled endomembrane compartments (Figure 1G). The fluorescence signal from the remaining nine fusion proteins had a non-informative distribution in the nucleus and the cytosol, similar to the distribution of a free GFP control (Figure 1H-P).

To explain the specific accumulation patterns observed, we examined the candidate effectors for subcellular targeting sequences. We focused on PST15391 and PST18447, because they showed specific, robust accumulation in nuclei (Figure 1D and 1F; Figure 2). Previous analysis failed to identify a nuclear-localisation signal (NLS) for PST15391 and PST18447 (Cantu *et al*., 2013). However, we noted that both carry NLS-like stretches of amino acids enriched in positively charged residues at the C-terminus and N-terminus of their mature forms, respectively (Figure 2A). Truncations lacking the NLS-like sequences accumulated mainly in the cytosol and only showed background accumulation in the nucleus, demonstrating that these regions are necessary for specific nuclear accumulation (Figure 2B). This set of experiments suggests that *P. striiformis* f sp *tritici* effectors use targeting sequences to traffic within plant cells.

**Figure 2.**
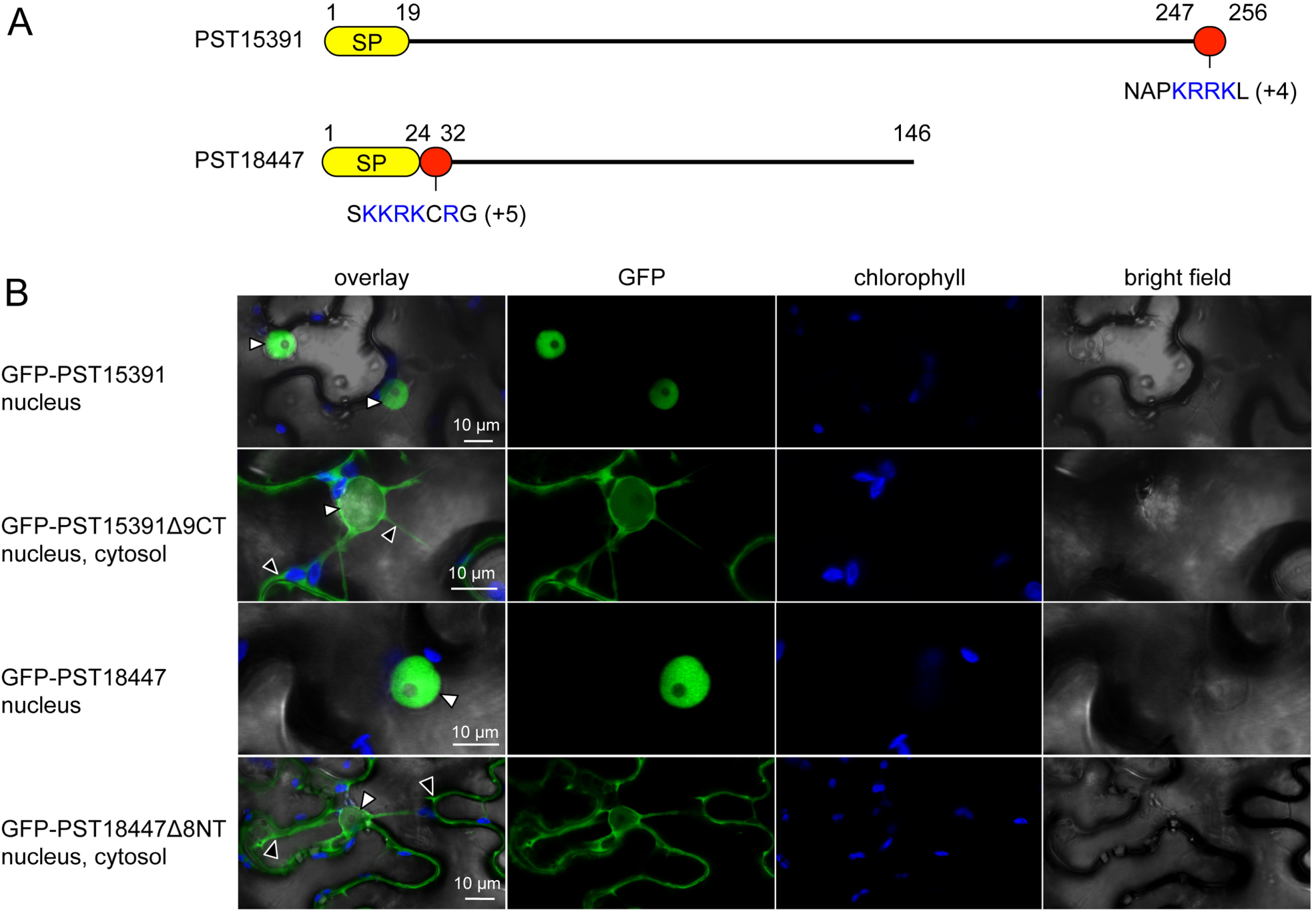
PST15391 and PST18447 carry functional nuclear-localisation signals. (A) Schematic representation of the protein primary structure of PST15391 and PST18447. Yellow: predicted signal peptide for secretion; red: amino acid sequence necessary for nuclear accumulation; blue: positively charged residues (net charge is indicated in parentheses). Numbers indicate amino acid positions. (B) Live-cell imaging of GFP-PST15391, GFP-PST15391A9CT, GFP-PST18447, and GFP-PST18447A8NT in *N. benthamiana* leaf cells. The cellular compartments in which the GFP signal accumulates are indicated. Proteins were transiently expressed in *N. benthamiana* leaf cells by agroinfiltration. Live-cell imaging was performed with a laser-scanning confocal microscope two days after infiltration. GFP and chlorophyll were excited at 488 nm. GFP (green) and chlorophyll (blue) fluorescence were collected at 505-525 nm and 680-700 nm, respectively. Images are single optical sections of 0.8 μm or maximal projections of up to 3 optical sections (max. z-stack of 2.4 μm). White arrowheads: nuclei; black arrowheads: cytosol.

### Six candidate effectors specifically and reliably associate with plant proteins

To gain further insight into the putative functions of the candidate effectors, we next aimed to identify the plant proteins they interact with *in planta* using anti-GFP coimmunoprecipitation/liquid chromatography-tandem mass spectrometry (coIP/MS) (Petre *et al*., 2015a). Using this approach, we identified 439 *N. benthamiana* proteins as potential interactors of the 16 candidate effectors (Table S2, Figure S3). A candidate effector associated with an average of 98 proteins, ranging from 20 to 236 (Figure 3A). Conversely, a plant protein associated with an average of 3.5 candidate effectors, ranging from 1 to 16 (Figure 3B). Given the high complexity of the dataset, we used a scoring method we previously developed to discriminate reliable and specific interactors (high score) from redundant and non-specific ones (low score) (Petre *et al*., 2015a). Scores ranged from 0.003 to 108, with an average value of 0.91 (Figure 3C). Eighteen proteins had a score of ≥ 3, and all specifically coimmunoprecipitated with a single candidate effector (Figure 3C, Figure 4). For instance, the protein with the highest score (108) was an enhancer of mRNA decapping protein 4 (NbEDC4) that specifically and robustly immunoprecipitated with PST02549 (Table S2).

**Figure 3.**
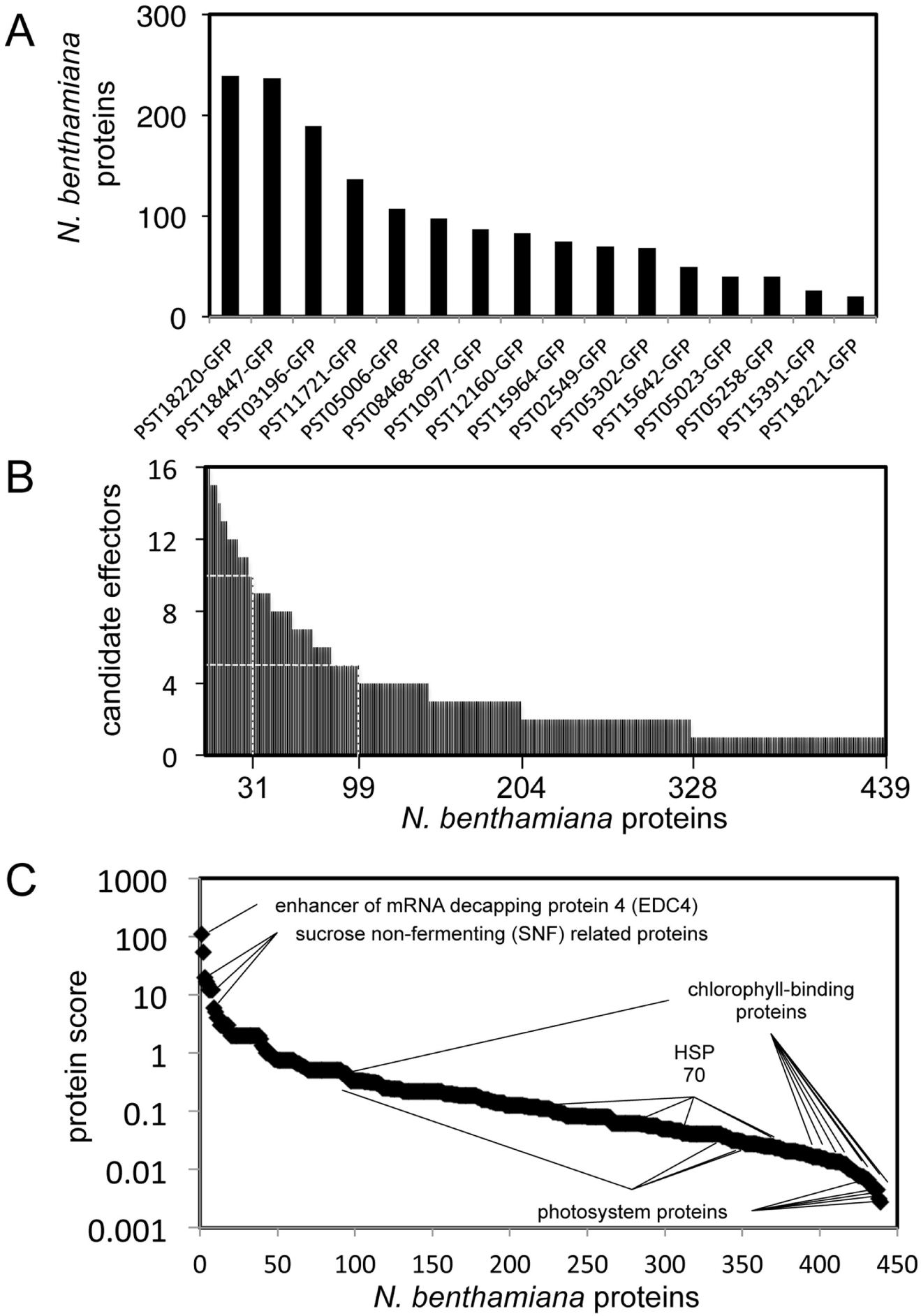
Candidate effectors associate with distinct plant protein complexes. (A) Number of *N. benthamiana* proteins associating with each candidate effector. Candidate effectors are arranged from left to right in descending order according to the number of interactors. (B) Number of candidate effectors associating with each *N. benthamiana* protein. The 439 interactors are arranged from left to right in descending order according to the number of associated candidate effectors. The X-axis legend indicates (from right to left) the number of *N. benthamiana* proteins that associated with at least one (439), two (328), three (204), five (99), and ten (31) candidate effectors. (C) For each *N. benthamiana* protein identified, we calculated a score following the formula “protein score = maximal peptide count/(redundancy)^2^”. The redundancy value was calculated by integrating the coIP/MS data from Petre *et al*., 2015a. Proteins are arranged from left to right in descending order based on their score. Selected proteins are indicated on the graph. Proteins were transiently expressed in *N. benthamiana* leaf cells by agroinfiltration. Total proteins were isolated two days after infiltration. Plant protein complexes associated with the candidate effector-GFP fusions were purified by anti-GFP coimmunoprecipitation, separated with SDS-PAGE, and digested with trypsin. Trypsin-digested peptides were processed by LC-MS/MS and collected peaks were used to search a database containing the predicted proteome of *N. benthamiana*. After filtering out contaminants and proteins supported by a single peptide, and clustering similar proteins, a total of 439 non-redundant protein interactors were retained. The full dataset used to draw these figures is shown in Table S2.

**Figure 4.**
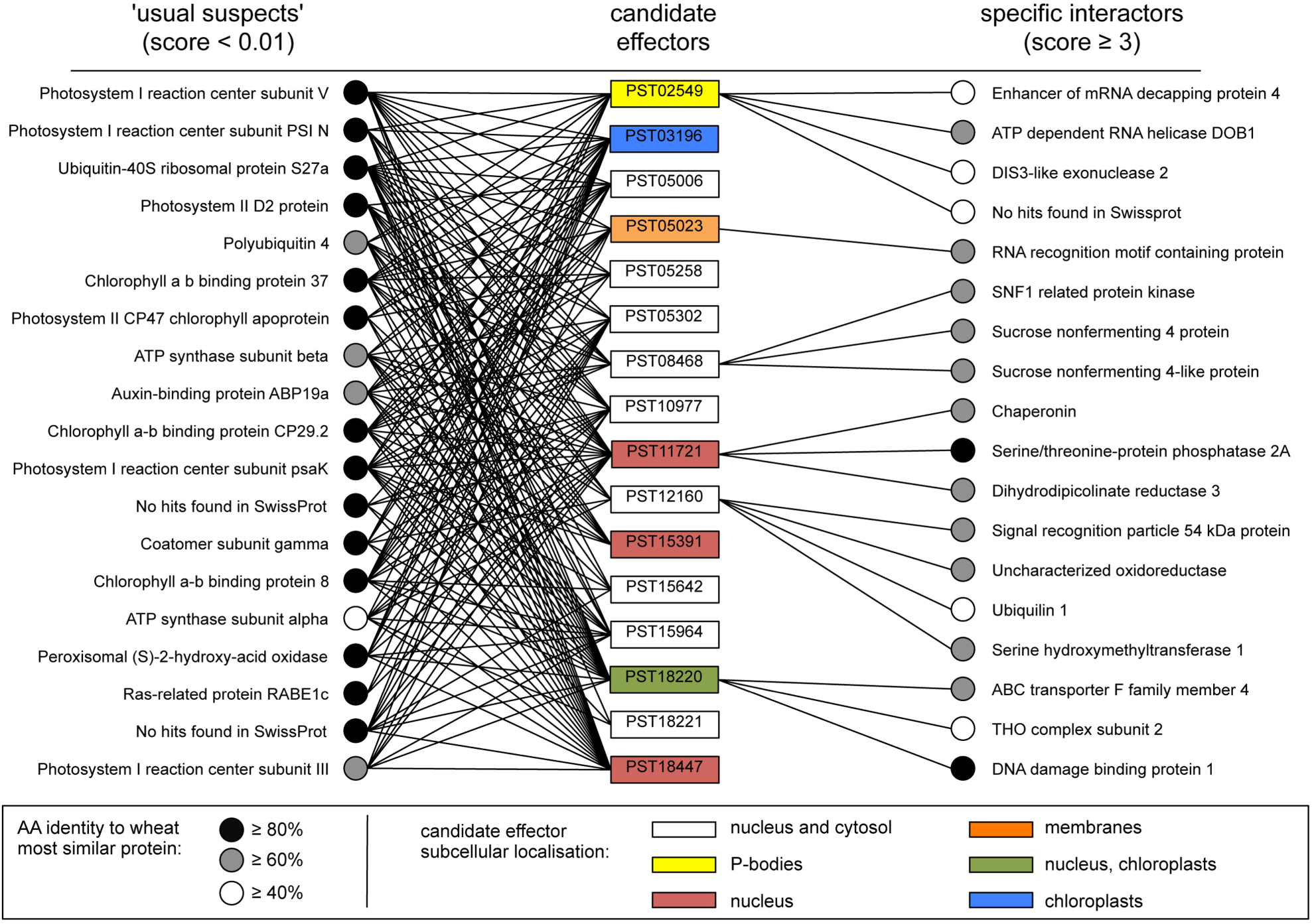
Nine candidate effectors have a specific subcellular localisation and/or a high-scoring plant protein interactor. The 16 candidate effectors used in this study are shown in the middle column. Colours indicate specific subcellular localisation. The 16 plant proteins with the lowest scores (≤ 0.01; termed ‘usual suspects’) and the 18 plant proteins with the highest scores (≥ 3; termed ‘specific interactors’) are shown on the left- and right-hand sides, respectively. Black lines indicate the association between a candidate effector and a plant protein as detected by coIP/MS. For each *N. benthamiana* protein, the most similar wheat protein was identified by protein sequence similarity searches against the predicted proteome of the bread wheat *Triticum aestivum* L. using the BLASTp algorithm.

### PST02549 associates with the wheat enhancer of mRNA decapping protein 4 (TaEDC4) in P-bodies

Our coIP/MS assays showed that PST02549 specifically associated with NbEDC4. To evaluate the biological significance of this association, we first identified and cloned the protein with the highest amino acid sequence similarity to NbEDC4 in bread wheat (*Triticum aestivum*), and named it TaEDC4. NbEDC4, TaEDC4, and *Arabidopsis thaliana* EDC4 (AtEDC4, also known as VARICOSE or VCS) are of a similar length (1203 to 1349 amino acids) and exhibit a pairwise amino acid sequence identity of between 42 and 46% (Figure 5A). The amino acid sequence identity between these proteins reaches 75% in the N-terminal WD40 domain (Figure 5B). Next, we expressed a TaEDc4-mCherry fusion in *N. benthamiana* leaf cells. Confocal microscopy revealed that TaEDC4-mCherry accumulated in cytosolic foci in addition to the cytosol. Since EDC4 is a component of P-bodies, we hypothesized that the foci we observed were P-bodies. To test this hypothesis, we co-expressed TaEDC4-mCherry with YFP-VCSc, a marker of P-bodies (Xu *et al*., 2006). Confocal microscopy revealed perfectly overlapping signals in cytosolic foci, confirming that TaEDC4 accumulates in P-bodies in *N. benthamiana* leaf cells (Figure 5C).

**Figure 5.**
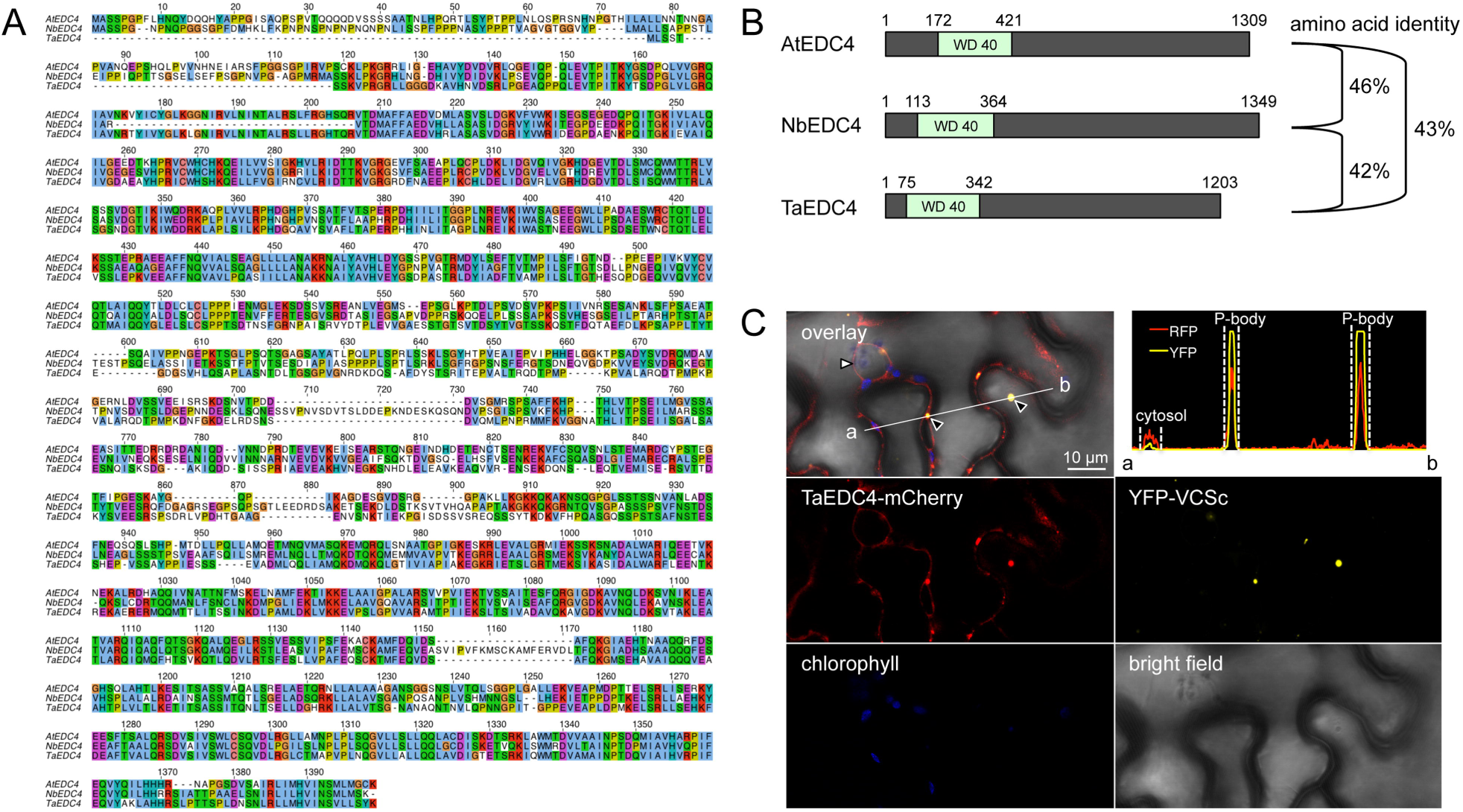
TaEDC4 accumulates in P-bodies. (A) Amino acid alignment of EDC4 of *Arabidopsis thaliana* (AtEDC4, AT3G13300.2), *Nicotiana benthamiana* (NbEDC4, NbS00023257g0003.1), and *Triticum aestivum* (TaEDC4, Traes_6DL_3FBA5B70E.1). Alignment was performed with ClustalX. Amino acid residues are colored according to the ClustalX scheme. (B) Schematic representation of the protein primary structure of AtEDC4, NbEDC4, and TaEDC4. Numbers indicate amino acid positions. The percentage of pairwise amino acid sequence identity is indicated to the right of the diagram. (C) Live-cell imaging of TaEDC4-mCherry and YFP-VCSc in *N. benthamiana* leaf cells. Images show a single optical section of 0.8 μm. Proteins were transiently expressed in *N. benthamiana* leaf cells by agroinfiltration. Live-cell imaging was performed with a laser-scanning confocal microscope with a sequential scanning mode two days after infiltration. The YFP was excited at 514 nm; mCherry and chlorophyll were excited at 561 nm. YFP (yellow), mCherry (red), and chlorophyll (blue) fluorescence were collected at 525-550 nm, 580-620 nm, and 680-700 nm, respectively. White arrowhead: nuclei; black arrowhead: P-bodies. The intensity plot in the top right corner shows YFP and mCherry (RFP) relative fluorescence signal intensity along the white line connecting points a and b in the overlay image.

To determine whether PST02549 and TaEDC4 associate *in planta*, we co-expressed PST02549-GFP and TaEDC4-mCherry fusion proteins in *N. benthamiana* leaf cells and performed anti-GFP coimmunoprecipitation followed by immunoblotting or sodium dodecyl sulphate polyacrylamide gel electrophoresis/Coomassie Brilliant Blue (SDS-PAGE/CBB) staining. Both the anti-mCherry immunoblot and the SDS-PAGE/CBB assays revealed a specific band signal matching the predicted size of TaEDC4-mCherry in protein complexes immunoprecipitated with PST02549-GFP, indicating a strong and robust association between the two proteins (Figure 6). As negative controls, we used three GFP and three mCherry fusion proteins available in the lab (see Materials and Methods for details); none of these control proteins associated with either PST02549 or TaEDC4. Confocal microscopy revealed that the fluorescence signals from PST02549-GFP and TaEDC4-mCherry perfectly overlapped in cytosolic foci, indicating co-accumulation in P-bodies (Figure 7A). From this set of experiments, we conclude that PST02539 and TaEDC4 specifically and robustly associate in P-bodies in *N. benthamiana* leaf cells.

**Figure 6.**
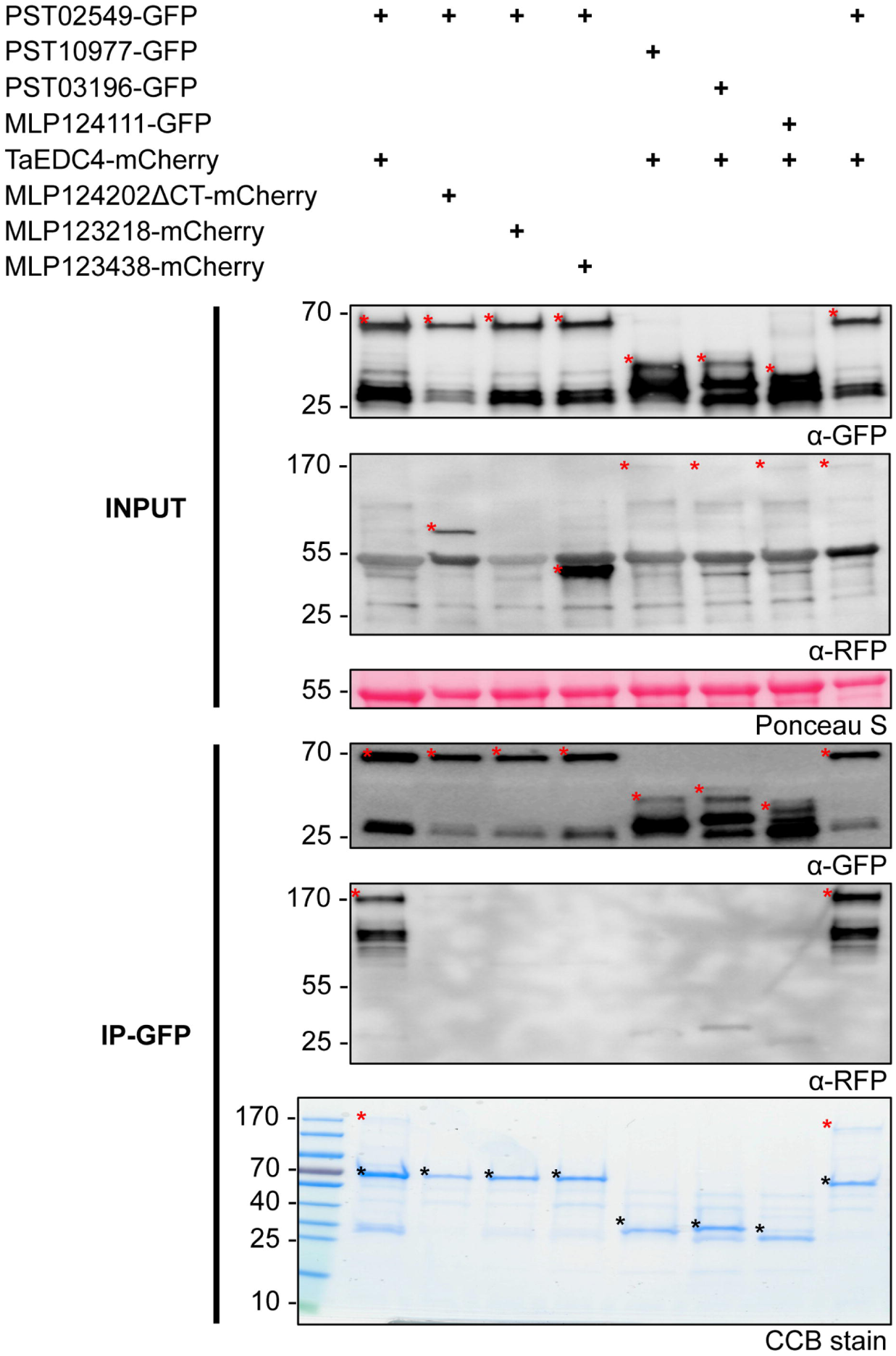
PST02549 associates with TaEDC4 *in planta*. Anti-green fluorescent protein (GFP) coimmunoprecipitation followed by immunoblot and sodium dodecyl sulphate-polyacrylamide gel electrophoresis/Coomassie Brilliant Blue (SDS-PAGE/CBB) analyses. Proteins were transiently expressed in *N. benthamiana* leaf cells by agroinfiltration. Total proteins were isolated two days after infiltration, and immediately used for anti-GFP immunoprecipitation. Immunoprecipitated protein mixtures were separated with SDS-PAGE. For direct protein visualization, the acrylamide gel was stained with CBB. For immunoblotting, proteins were electrotransferred onto polyvinylidene fluoride (PVDF) membranes. Immunodetection was performed with anti-GFP or anti-redu fluorescent protein (RFP) antibodies, and immunoblots were revealed with a chemiluminescent imager. Ponceau S staining of the PVDF membrane was used as a loading and transfer control. Theoretical protein size is indicated in parentheses in kilodalton (kDa) for each fusion protein. Numbers to the left of the blot and gel images indicate protein size in kDa. In the immunoblot images, red asterisks indicate specific protein bands. In the gel image, asterisks indicate specific protein bands (red: TaEDC4-mCherry; black: GFP fusions); the PageRuler ladder is shown to the left of the image. IP: immunoprecipitation. In the IP-GFP/ α-RFP blot, note that the weak band signals observed on the right side between 25 and 40 kDa are due to non-specific background detection of abundant GFP fusions by the anti-RFP antibodies.

### Co-expression of PST02549 and TaEDC4 increases the size of P-bodies

During confocal microscopy assays of *N. benthamiana* leaf cells co-expressing PST02549-GFP and TaEDC4-mCherry, we noted that the P-bodies appeared larger than usual (Figure 7A). To quantify this phenomenon, we measured the diameter of P-bodies from confocal microscopy images. When PST02549-GFP was co-expressed with TaEDC4-mCherry or an untagged version of TaEDC4, the average diameters of the P-bodies were 4.5 ± 3.5 μm and 4.9 ± 2.2 μm, respectively (Figure 7B, Table S4). By contrast, when PST02549-GFP and TaEDC4-mCherry were expressed independently and/or with other control proteins, the average diameter of a P-body was 1.3 ± 0.6 μm. CoIP/MS assays confirmed the presence of the untagged TaEDC4, as well as the presence of the endogenous NbEDC4, in complex with PST02549 (Table S5). None of the negative controls we tested co-localised with PST02549-GFP or TaEDC4-mCherry or triggered the formation of large P-bodies (Figure 7B and C, Table S4). We conclude that co-expression of PST02549 and TaEDC4 specifically increases the size of P-bodies.

**Figure 7.**
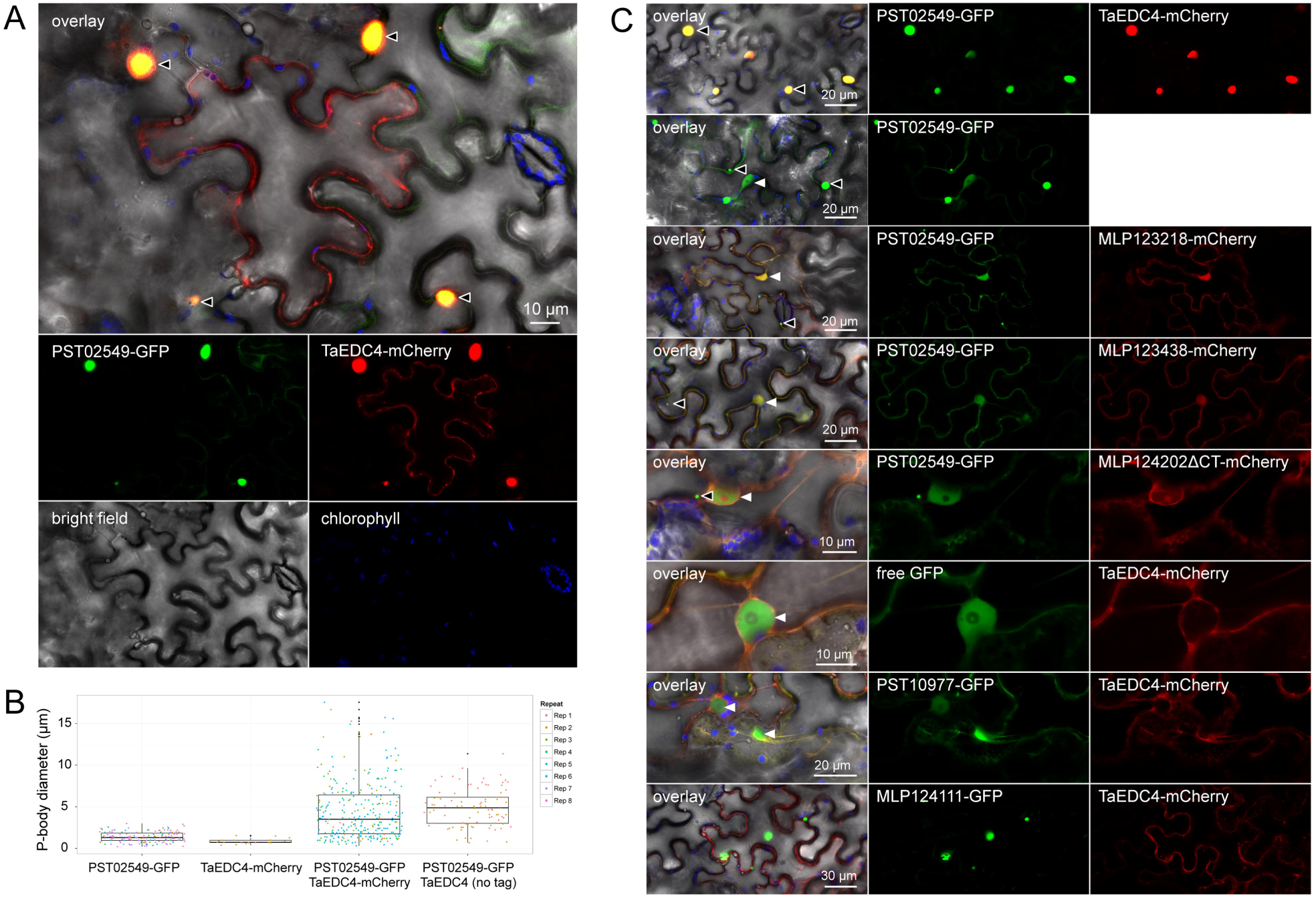
PST02549 and TaEDC4 co-accumulate in large P-bodies. (A) Live-cell imaging of PST02549-GFP and TaEDC4-mCherry in *N. benthamiana* leaf cells. Images show a single optical section of 0.8 µm. The white asterisk indicates a pavement cell expressing only the TaEDC4-mCherry fusion, in which no large P-body was detected. (B) Categorical scatterplots showing the diameter of P-bodies labelled by PST02549-GFP and/or TaEDC4-mCherry in leaf cells. Boxes depict the interquartile range and the median, vertical bars indicate the first and fourth quartile range, and outlier data points are depicted in black. P-body diameters were measured from laser scanning confocal microscope images acquired through two to eight independent agroinfiltration assays. The different colours correspond to independent observations (repeats). The following numbers of P-bodies were scored: PST02549-GFP (n=150); TaEDC4-mCherry (n=20), PST02549-GFP/TaEDC4-mCherry (n=303), PST02549-GFP/TaEDC4 (n=96). For treatments ‘PST02549-GFP’ and ‘TaEDC4-mCherry’, the fusion proteins were expressed alone or with additional control fusion proteins (see Table S4 for raw data). (C) Live-cell imaging of various GFP and mCherry fusion proteins in *N. benthamiana* leaf cells. Images present a single optical section of 0.8 µm of a maximal projection of up to 6 optical sections (max. z-stack of 4.8 µm). Overlay images merge GFP, mCherry, chlorophyll, and bright field signals. Note that for the PST02549-GFP/TaEDC4, TaEDC4 was untagged and the mCherry fluorescence signal was not recorded. For (A) and (C), proteins were transiently expressed in *N. benthamiana* leaf cells by agroinfiltration. Live-cell imaging was performed with a laser-scanning confocal microscope with a sequential scanning mode two days after infiltration. GFP and the chlorophyll were excited at 488 nm; the mCherry was excited at 561 nm. GFP (green), mCherry (red), and chlorophyll (blue) fluorescence were collected at 505-525 nm, 580-620 nm and 680-700 nm, respectively. Black arrowheads indicate P-bodies. White arrowheads: nuclei. Note that the large protein aggregates formed by MLP124111-GFP do not show any TaEDC4-mCherry signal.

## DISCUSSION

In this study, we found that PST02549 accumulates in plant cell P-bodies and associates with a P-body-derived protein. This observation suggests that an effector that targets plant P-bodies has evolved in *P. striiformis* f sp *tritici*. To our knowledge, a connection between filamentous plant pathogens and P-bodies has not previously been established.

How would a pathogen benefit from manipulating host P-bodies? Some plant pathogen effectors target components of the host RNA silencing machinery (Weiberg *et al*., 2013; Qiao *et al*., 2015, Spanu 2015). In *A. thaliana*, two recent reports have connected P-bodies, or P-body-resident proteins, with plant immune responses (Maldonado-Bonilla et al., 2014; Roux *et al*., 2015). Therefore, pathogen effectors may target P-bodies to manipulate RNA metabolism and/or suppress immune responses. Interestingly, pathogenic bacteria and viruses that infect mammals are known to alter the structure and function of host P-bodies (Ariumi *et al*., 2011; Eulalio *et al*., 2011; Perez-Vilaro *et al*., 2012). Therefore, diverse parasites of eukaryotes have evolved to target host P-bodies. Further mechanistic investigations of the pathogen effector/P-body interplay should reveal the biological significance of this phenomenon.

We observed an increase in P-body size upon co-expression of PST02549 and TaEDC4. The depletion or overexpression of P-body components is known to modify P-body integrity, which can lead to an increase in size (Eulalio *et al*., 2007). It is therefore possible that the increase in P-body size observed in our study is due to over-accumulation of P-body-resident proteins such as PST02549 or TaEDC4. However, we observed this phenomenon only when the two proteins co-accumulated, indicating that both are required to increase P-body size. The biological significance of the association between PST02549 and TaEDC4 as well as the increase in P-body size remain to be further investigated in wheat.

The pipeline we used in this study allowed us to retrieve informative data for more than 50% of the candidate effectors we tested. We recently obtained informative data for 40% of a set of candidate effectors from another rust species (Petre *et al*., 2015a). Thus, *N. benthamiana* is a valuable heterologous system for fast-forward effectoromic analysis of plant pathogens, regardless of their host plant.

We identified plant interactors of candidate effectors, some of which may represent *bone fide* effector targets. Growing evidence suggests that during evolution domains from effector targets have been incorporated into immune receptors such as nucleotide binding-leucine rich repeat (NB-LRR, also referred to as NLR) proteins to become ‘sensor domains’ that mediate recognition of specific effectors (Cesari *et al*., 2014; Wu *et al*., 2015; Sarris *et al*., 2015). A recent genome-wide analysis predicted many NLR gene models in which protein domains that differ from typical NLR domains have been incorporated (Sarris *et al*., in review). Interestingly, six of the eighteen top scoring effector interactors identified in our study carry a protein domain that is predicted to be integrated into a plant NLR, including the WD40 protein domain of EDC4 (Table S2). Therefore, our predicted host targets can be a valuable source of new ‘baits’ for engineering NLR genes with sensor domains.

## MATERIALS AND METHODS

### *In silico* analyses

Predicted protein sequences were retrieved from the following sources: *P. striiformis f sp tritici* (http://yellowrust.com/; Cantu *et al*., 2013), *N. benthamiana* (http://solgenomics.net/; Bombarely *et al*., 2012), and *T. aestivum* (http://www.plantgdb.org/TaGDB/). Protein sequence analysis was performed with ClustalX and Jalview programmes. Homology searches were performed with the BLAST+ programme. The most stringent criterion for selection of the candidate effectors was transcript enrichment in purified haustoria compared to infected tissues (Cantu *et al*., 2013). The set of candidate effectors selected via the pipeline was manually analysed to remove redundant family members. PST05258 and PST15391 from Tribe 54, as well as PST18220 and PST18221 from Tribe 238, were both retained due to high levels of polymorphism (Cantu *et al*., 2013).

### Cloning procedures and plasmids

The open reading frame (ORF) encoding the mature form (i.e. without the signal peptide) of *P. striiformis* f sp *tritici* small-secreted proteins or the full-length of *T. aestivum* EDC4 (Traes_6DL_3FBA5B70E.1) were amplified by polymerase chain reaction (PCR) using cDNA isolated from wheat leaves 14 days after inoculation with a virulent isolate of *P. striiformis* f sp *tritici* (isolate PST-08/21; Cantu *et al*., 2013), or were obtained through gene synthesis (Genewiz, London, UK), with codon optimization for plant expression and removal of internal Bbsl and Bsal restriction sites. Primers and synthetic genes were designed to be compatible with the suite of Golden Gate vectors, as previously described (Petre *et al*., 2015a; Table S1). Truncated versions of PST15391 and PST18447 were obtained by PCR cloning. All PCR-generated DNA fragments were verified by sequencing after cloning into level 0 Golden Gate vectors. Plasmids were multiplied and conserved in *Escherichia coli* (Subcloning Efficiency DH5α Competent Cells; Invitrogen, Carlsbad, California, USA) as previously described (Petre *et al*., 2015a). The fusion proteins built with candidate effectors from the poplar rust fungus *Melampsora larici-populina* (MLP124111, MLP123218, MLP123438 and MLP124202CT) were obtained in previous studies (Petre *et al*., 2015a, Petre *et al*., 2015b, Petre *et al*., unpublished) and were used as negative controls in coIP and confocal microscopy assays.

### Transient protein expression in *N. benthamiana* leaf cells

*Agrobacterium tumefaciens* (electrocompetent strain GV3101) was used to deliver T-DNA constructs in leaf cells of three- to four-week-old *N. benthamiana* plants, following the agroinfiltration method previously described (Petre *et al*., 2015a). The leaves were collected two days after infiltration for further protein isolation or microscopy.

### Live-cell imaging by laser-scanning confocal microscopy

Confocal microscopy was performed as previously reported (Petre *et al*., 2015a) with a Leica DM6000B/TCS SP5 laser-scanning confocal microscope (Leica Microsystems, Bucks, UK), using 10x (air) and 63x (water immersion) objectives. Each construct gave a similar localisation pattern in at least three independent observations. Image analysis was performed using the Fiji plugin of Image J 2.0.0 (http://fiii.sc/Fiii). To quantify the diameter of P-bodies, the ‘measure’ tool of Fiji was used to measure manually-drawn lines matching the apparent diameter of P-bodies in all the single optical section confocal images acquired in the course of this project. Categorical scatterplots were generated with R, using the ggplot2 package and an in-house developed script (Text S1).

### Protein isolation and immunoblot analyses

Frozen leaves were ground to a powder using a mortar and pestle. Total proteins were extracted as previously described (Petre *et al*., 2015a). Ten microliters of isolated protein was separated on a 15% SDS-PAGE gel, and the protein content was estimated by Coomassie Brilliant Blue (CBB) staining. Immunoblot analysis was performed as previously described (Petre *et al*., 2015a), using GFP (B2):sc-9996 HRP-conjugated antibody (Santa-Cruz Biotechnology), rat anti-RFP 5F8 antibody (Chromotek, Munich, Germany) and a HRP-conjugated anti-rat antibody.

### Coimmunoprecipitation and LC-MS/MS analyses

Coimmunoprecipitation procedures were performed as reported by Win and colleagues (2011), with the adaptation described in Petre *et al*., 2015a, using GFP_Trap_A beads (Chromotek, Munich, Germany). GFP and mCherry fusion proteins (Petre *et al*., 2015a, Petre *et al*., 2015b, Petre *et al*., unpublished) selected based on their ability to generate cytosolic aggregates, their similarity to tested proteins, or their ability to associate with a high number of proteins (i.e. their ‘stickyness’) in coIP assays were used as negative controls. Sample preparation, liquid chromatography / tandem mass spectrometry (LC-MS/MS) and data analyses were performed as described in Petre *et al*., 2015a, using a hybrid mass spectrometer LTQ-Orbitrap XL (ThermoFisher Scientific, Carlsbad, California, USA) and a nanoflow-UHPLC system (NanoAcquity Waters Corp., Burnsville, Minnesota, USA). LC-MS/MS data were processed and scored as previously described (Petre *et al*., 2015a; Table S2).

## ACKNOWLEDGMENTS

We thank the Norwich Rust Group for discussions, Dr. Vanessa Segovia (Norwich, UK) for providing material, and support teams at The Sainsbury Laboratory and The John Innes Centre. BP was supported by an INRA Contrat Jeune Scientifique (CJS), by the European Union, in the framework of the Marie-Curie FP7 COFUND People Programme, through the award of an AgreenSkills’ fellowship (under grant agreement n° 267196). DGOS was supported by a Leverhulme early career fellowship and a fellowship in computational biology at TGAC, in partnership with the John Innes Centre, and strategically supported by BBSRC. CL is supported by an INRA CJS. BP, CL, and SD are supported by the French National Research Agency through the Labex ARBRE (ANR-12-LABXARBRE-01) and the Young Scientist Grant POPRUST (ANR-2010-JCJC-1709-01). KVK is strategically supported by the BBSRC and the Gatsby Charitable Foundation. Research at The Sainsbury Laboratory is supported by the Gatsby Charitable Foundation and the BBSRC.

---

**Figure S1.**
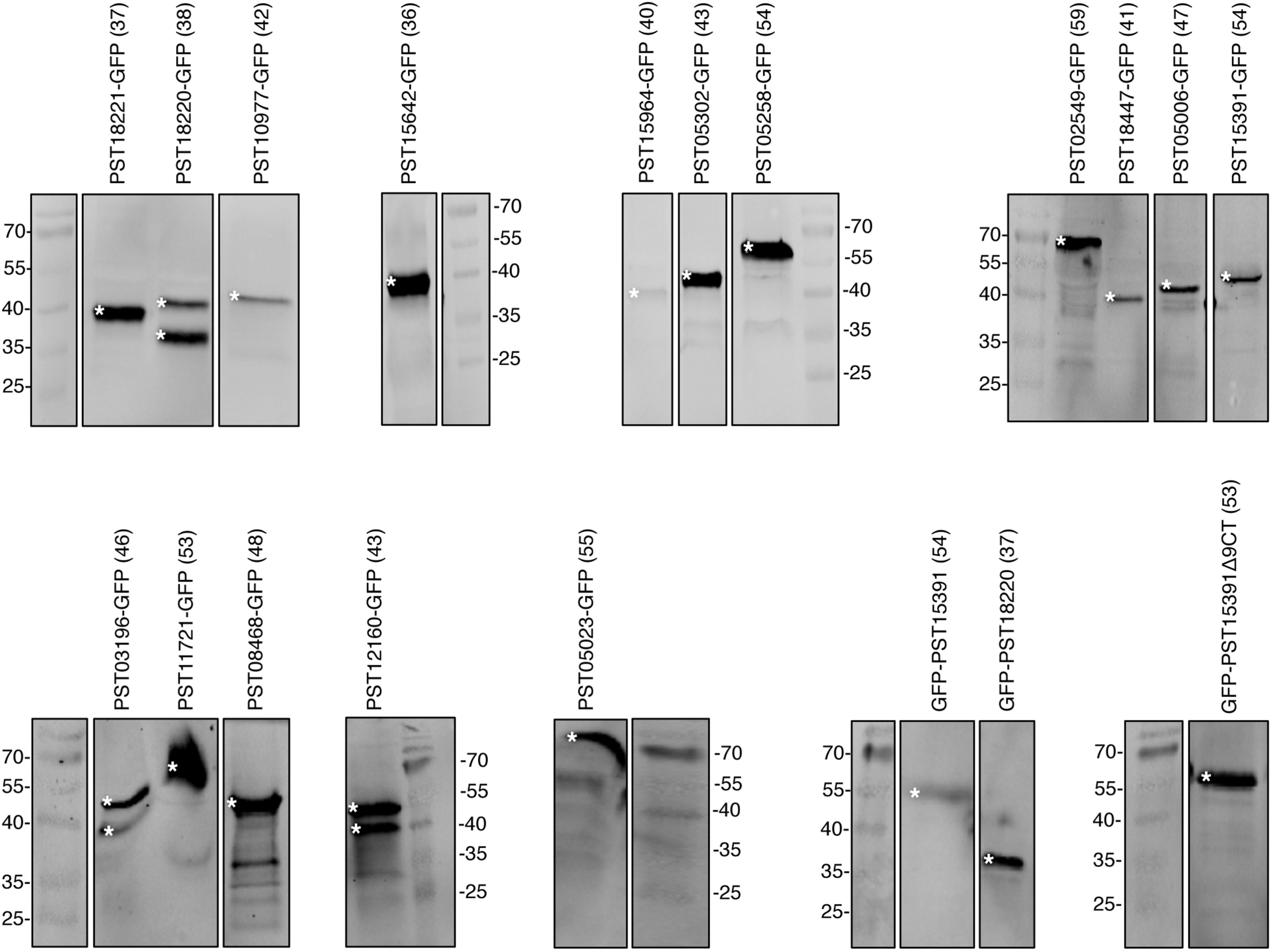
Immunoblots confirm the accumulation of the fusion proteins in *N. benthamiana* leaf cells. Proteins were transiently expressed in *N. benthamiana* leaf cells by agroinfiltration. Total proteins were extracted two days after infiltration by grinding leaves in liquid nitrogen and immediately extracting, reducing and denaturing proteins from the leaf powder. Proteins were separated on 15% SDS-PAGE gels and transferred onto a nitrocellulose membrane. Primary and secondary immune detection were performed with rabbit anti-GFP and goat anti-rabbit antibodies, respectively. Images originating from the same membrane and processed at the same time are grouped together. Blots were cropped to remove lanes previously published elsewhere (Petre *et al*., 2015a). Secondary antibodies and PageRuler signals were detected simultaneously using an infrared imager. The theoretical size of each fusion protein is indicated in kDa in parentheses. White asterisks indicate specific protein bands.

**Figure S2.**
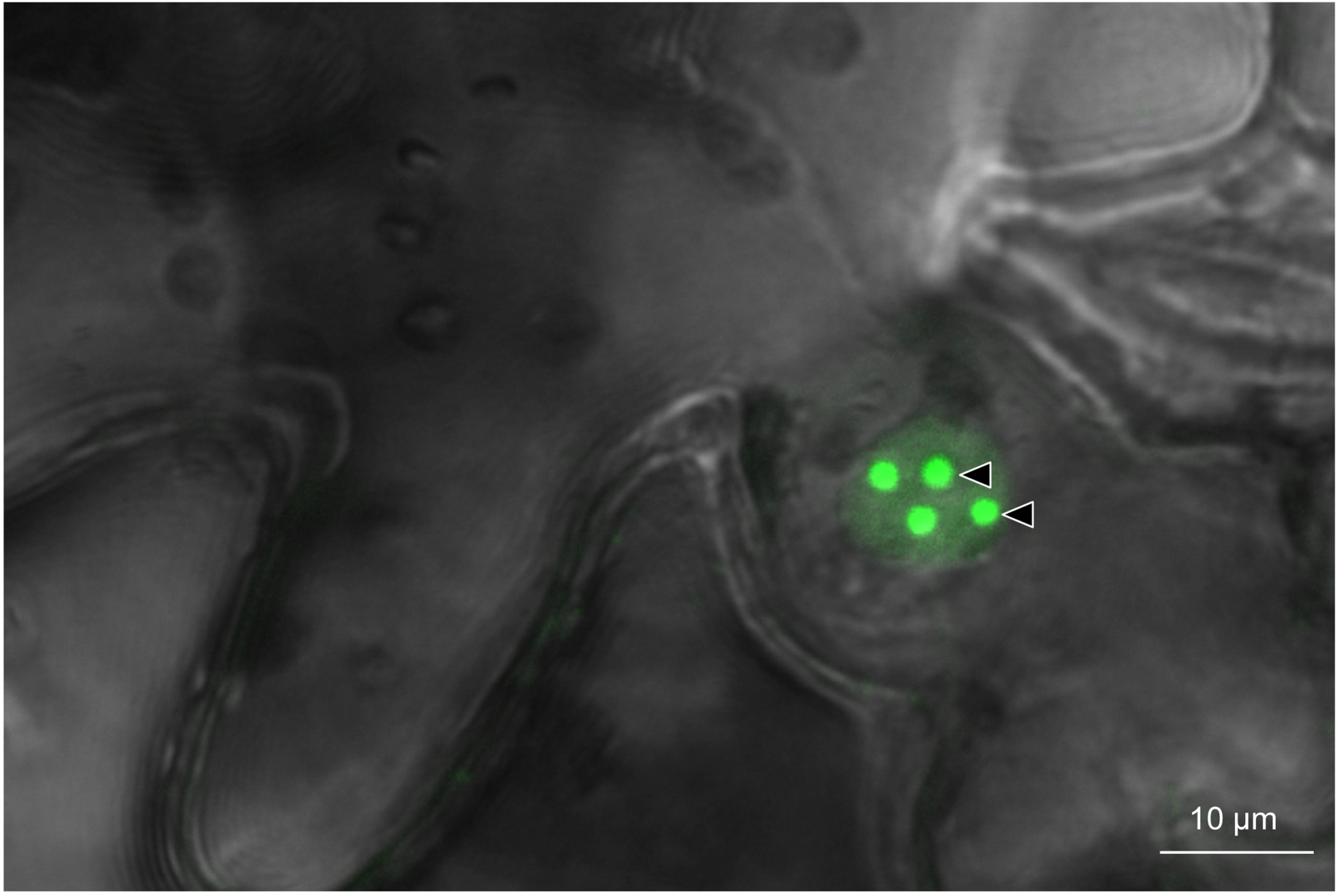
PST11721-GFP labels nuclei foci. Live-cell imaging of PST11721-GFP in *N. benthamiana* leaf cells. Proteins were transiently expressed in *N. benthamiana* leaf cells by agroinfiltration. Live-cell imaging was performed with a laser-scanning confocal microscope two days after infiltration. The GFP was excited at 488 nm. GFP (green) fluorescence was collected at 505-550 nm. The image is a single optical section of 0.8 µm, showing an overlay of the GFP and bright field channels. The black arrowheads indicate GFP-labelled nuclear foci.

**Figure S3.**
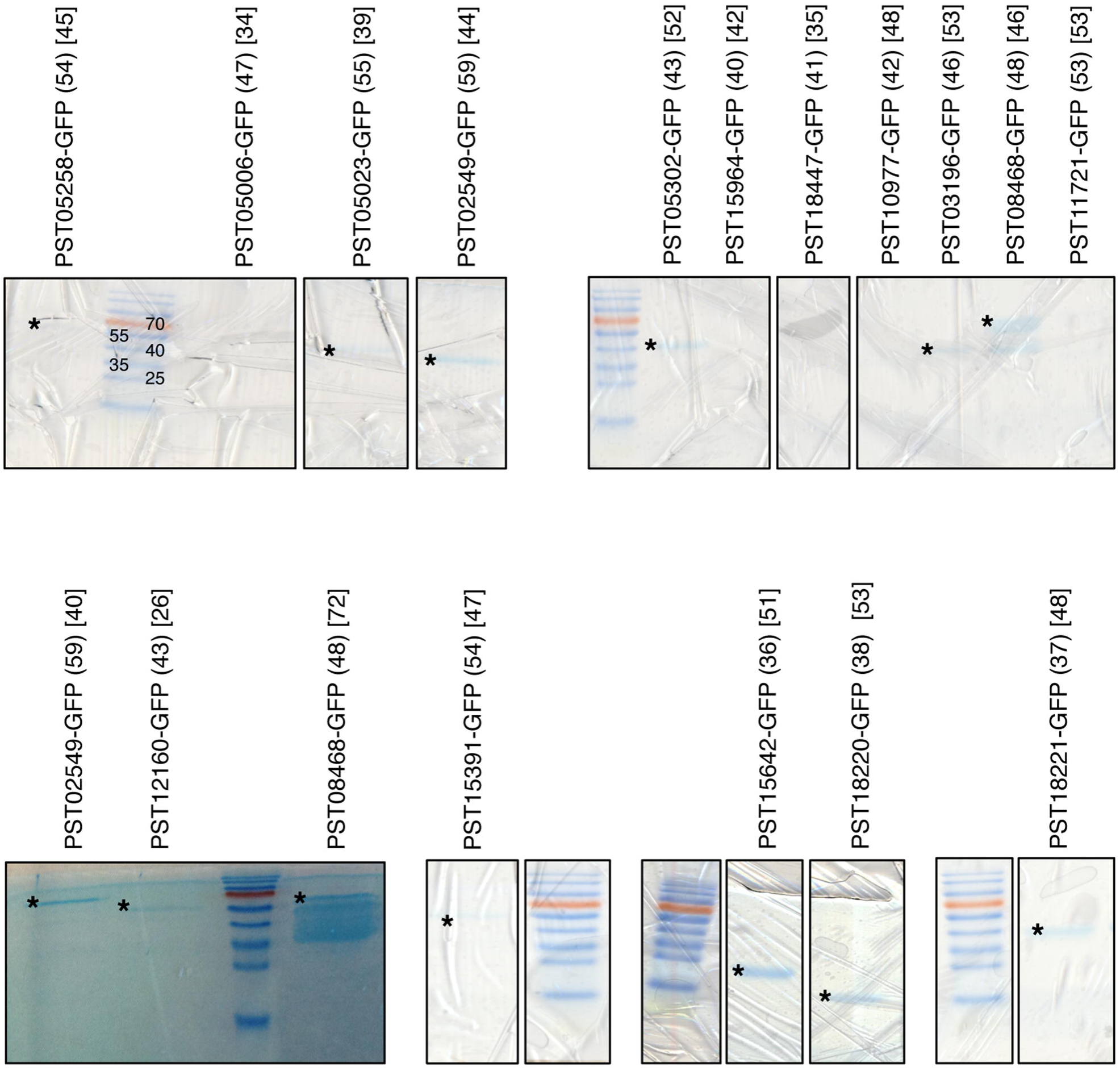
*In planta* coimmunoprecipitation efficiently purifies fusion proteins. Protein mixtures isolated by anti-GFP immunoprecipitation were reduced and denatured in a Laemmli buffer. Proteins were separated with SDS-PAGE and stained with Coomassie Brilliant Blue. Trypsin-digested peptides were processed by LC-MS/MS and collected peaks were used to search a database containing the GFP sequence. The theoretical size of each fusion protein is indicated in parentheses in kilodalton (kDa). The number of peptides identified by LC-MS/MS and matching the GFP is indicated for each fusion protein between brackets. The size of the PageRuler ladder bands is indicated in kDa. Images originating from the same gel are grouped together. Black asterisks indicate detectable and specific protein bands.

TEXT S1. R script to generate categorical scatterplots

**TABLE S1.** Cloning and protein details

**TABLE S2.** The *P. striiformis* f sp *tritici* candidate effector interactome

**TABLE S3.** Overlap between CoIp/MS replicates

**TABLE S4.** P-body diameter values

**TABLE S5.** Detection of TaEDC4 in *N. benthamiana* leaves by coIP/MS

## REFERENCES

Ariumi Y, Kuroki M, Kushima Y, Osugi K, Hijikata M, Maki M, Ikeda M, Kato N. 2011. Hepatitis C virus hijacks P-body and stress granule components around lipid droplets. J Virol 85: 6882–92

Beddow JM, Pardey PG, Chai Y, Hurley TM, Kriticos DJ, Braun H-J, Park RF, Cuddy WS, Yonow T. 2015. Research investment implications of shifts in the global geography of wheat stripe rust. Nat Plants 1: 15132

Bombarely A, Rosli HG, Vrebalov J, Moffett P, Mueller LA, Martin GB. 2012. A draft genome sequence of *Nicotiana benthamiana* to enhance molecular plant-microbe biology research. Mol Plant Microbe Interact 25: 1523–30

Cantu D, Segovia V, MacLean D, Bayles R, Chen X, Kamoun S, Dubcovsky J, Saunders DG, Uauy C. 2013. Genome analyses of the wheat yellow (stripe) rust pathogen *Puccinia striiformis* f. sp. *tritici* reveal polymorphic and haustorial expressed secreted proteins as candidate effectors. BMC Genomics 14: 270

Cantu D, Govindarajulu M, Kozik A, Wang M, Chen X, Kojima KK, Jurka J, Michelmore RW, Dubcovsky J. 2011. Next generation sequencing provides rapid access to the genome of *Puccinia striiformis* f. sp. *tritici*, the causal agent of wheat stripe rust. PLoS One 6:e24230

Cesari S, Bernoux M, Moncuquet P, Kroj T, Dodds PN. 2014. A novel conserved mechanism for plant NLR protein pairs: the “integrated decoy” hypothesis. Front Plant Sci 5: 606

Chen W, Wellings C, Chen X, Kang Z, Liu T. 2014. Wheat stripe (yellow) rust caused by *Puccinia striiformis* f. sp. *tritici*. Mol Plant Pathol 15: 433–46

Dangl JLl, Horvath DM, Staskawicz BJ. 2013. Pivoting the plant immune system from dissection to deployment. Science 341: 746–51

Dean R, Van Kan JA, Pretorius ZA, Hammond-Kosack KE, Di Pietro A, Spanu PD, Rudd JJ, Dickman M, Kahmann R, Ellis J, Foster GD. 2012. The Top 10 fungal pathogens in molecular plant pathology. Mol Plant Pathol 13: 414–30

Dodds PN, Rathjen JP. 2010. Plant immunity: towards an integrated view of plant-pathogen interactions. Nat Rev Genet 11: 539–48

Eulalio A, Fröhlich KS, Mano M, Giacca M, Vogel J. 2011. A candidate approach implicates the secreted Salmonella effector protein SpvB in P-body disassembly. PLoS One 6:e17296.

Eulalio A, Behm-Ansmant I, Izaurralde E. 2007. P bodies: at the crossroads of post-transcriptional pathways. Nat Rev Mol Cell Biol 8: 9–22

Garnica DP, Upadhyaya NM, Dodds PN, Rathjen JP. 2013. Strategies for Wheat Stripe Rust Pathogenicity Identified by Transcriptome Sequencing. PLoS One 8:e67150.

Goodin MM, Zaitlin D, Naidu RA, Lommel SA. 2008. *Nicotiana benthamiana:* its history and future as a model for plant-pathogen interactions. Mol Plant Microbe Interact 21: 1015–26

Hubbard A, Lewis CM, Yoshida K, Ramirez-Gonzalez RH, de Vallavieille-Pope C, Thomas J, Kamoun S, Bayles R, Uauy C, Saunders DG. 2015. Field pathogenomics reveals the emergence of a diverse wheat yellow rust population. Genome Biol 16: 23

Hogenhout SA, Van der Hoorn RA, Terauchi R, Kamoun S. 2009. Emerging concepts in effector biology of plant-associated organisms. Mol Plant Microbe Interact 22: 115–22

Maldonado-Bonilla LD, Eschen-Lippold L, Gago-Zachert S, Tabassum N, Bauer N, Scheel D, Lee J. 2014. The Arabidopsis tandem zinc finger 9 protein binds RNA and mediates pathogen-associated molecular pattern-triggered immune responses. Plant Cell Physiol 55: 412–25

Nemri A, Saunders DG, Anderson C, Upadhyaya NM, Win J, Lawrence GJ, Jones DA, Kamoun S, Ellis JG, Dodds PN. 2014. The genome sequence and effector complement of the flax rust pathogen Melampsora lini. Front Plant Sci 5: 98.

Pais M, Win J, Yoshida K, Etherington GJ, Cano LM, Raffaele S, Banfield MJ, Jones A, Kamoun S, Saunders DG. 2013. From pathogen genomes to host plant processes: the power of plant parasitic oomycetes. Genome Biol 14: 211

Pérez-Vilaró G, Fernández-Carrillo C, Mensa L, Miquel R, Sanjuan X, Forns X, Pérez-del-Pulgar S, Díez J. 2015. Hepatitis C virus infection inhibits P-body granule formation in human livers. J Hepatol 62: 785–90

Petre B, Saunders DG, Sklenar J, Lorrain C, Win J, Duplessis S, Kamoun S. 2015a. Candidate Effector Proteins of the Rust Pathogen Melampsora larici-populina Target Diverse Plant Cell Compartments. Mol Plant Microbe Interact 28: 689–700

Petre B, Lorrain C, Saunders DG, Win J, Sklenar J, Duplessis S, Kamoun S. 2015b. Rust fungal effectors mimic host transit peptides to translocate into chloroplasts. Cell Microbiol doi: 10.1111/cmi.12530

Petre B, Joly DL, Duplessis S. 2014. Effector proteins of rust fungi. Front Plant Sci 5: 416

Petre B, Kamoun S. 2014. How do filamentous pathogens deliver effector proteins into plant cells? PLoS Biol 12:e1001801

Qiao Y, Shi J, Zhai Y, Hou Y, Ma W. 2015. *Phytophthora* effector targets a novel component of small RNA pathway in plants to promote infection. Proc Natl Acad Sci U S A 112: 5850–5

Reineke LC, Lloyd RE. 2013. Diversion of stress granules and P-bodies during viral infection. Virology 436: 255–67

Roux ME, Rasmussen MW, Palma K, Lolle S, Regué AM, Bethke G, Glazebrook J, Zhang W, Sieburth L, Larsen MR, Mundy J, Petersen M. 2015. The mRNA decay factor PAT1 functions in a pathway including MAP kinase 4 and immune receptor SUMM2. EMBO J 34: 593–608

Sarris PF, Jones JD. 2015. Plant immune receptors mimic pathogen virulence targets. Oncotarget 6: 16824–5

Spanu PD. 2015. RNA-protein interactions in plant disease: hackers at the dinner table. New Phytol 207: 991–5

Steffens A, Bräutigam A, Jakoby M, Hülskamp M. 2015. The BEACH Domain Protein SPIRRIG Is Essential for Arabidopsis Salt Stress Tolerance and Functions as a Regulator of Transcript Stabilization and Localization. PLoS Biol 13:e1002188

Upadhyaya NM, Mago R, Staskawicz BJ, Ayliffe MA, Ellis JG, Dodds PN. 2014. A Bacterial Type III Secretion Assay for Delivery of Fungal Effector Proteins into Wheat. Mol Plant Microbe Interact 27: 255–264

Weiberg A, Wang M, Bellinger M, Jin H. 2014. Small RNAs: a new paradigm in plant-microbe interactions. Annu Rev Phytopathol 52: 495–516

Win J, Chaparro-Garcia A, Belhaj K, Saunders DG, Yoshida K, Dong S, Schornack S, Zipfel C, Robatzek S, Hogenhout SA, Kamoun S. 2012. Effector biology of plant-associated organisms: concepts and perspectives. Cold Spring Harb Symp Quant Biol 77: 235–47

Wu CH, Krasileva KV, Banfield MJ, Terauchi R, Kamoun S. 2015. The “sensor domains” of plant NLR proteins: more than decoys? Front Plant Sci 6: 134

Xu J, Chua NH. 2009. *Arabidopsis* decapping 5 is required for mRNA decapping, P-body formation, and translational repression during postembryonic development. Plant Cell 21: 3270–9

Xu J, Yang JY, Niu QW, Chua NH. 2006. *Arabidopsis* DCP2, DCP1, and VARICOSE form a decapping complex required for postembryonic development. Plant Cell 18: 3386–98

Zheng W, Huang L, Huang J, Wang X, Chen X, Zhao J, Guo J, Zhuang H, Qiu C, Liu J, Liu H, Huang X, Pei G, Zhan G, Tang C, Cheng Y, Liu M, Zhang J, Zhao Z, Zhang S, Han Q, Han D, Zhang H, Zhao J, Gao X, Wang J, Ni P, Dong W, Yang L, Yang H, Xu JR, Zhang G, Kang Z. 2013. High genome heterozygosity and endemic genetic recombination in the wheat stripe rust fungus. Nat Commun 4: 2673

